# NTCmatch: R package for searching synonyms of plants listed under list of Normally Traded as Commodities

**DOI:** 10.1101/2025.06.03.657558

**Authors:** Purnendu Paul, Prakash Pradhan

## Abstract

Equitable sharing of benefits from the commercial use of biological resources is a cornerstone of the Convention on Biological Diversity (CBD). This principle is pivotal in advancing the other two CBD objectives: conserving biological diversity and its sustainable use. India, as a CBD signatory, enacted the Biological Diversity Act in 2002 (amended in 2023) and established a three-tiered implementation system involving the National Biodiversity Authority (NBA), State Biodiversity Boards (SBBs), and Union Territory Biodiversity Councils (UTBCs). A critical regulatory framework within this system is the Access and Benefit Sharing (ABS) mechanism, guided by the Nagoya Protocol and NBA guidelines. This study addresses challenges in the identification and processing of biological resources listed as Normally Traded as Commodities (NTC), particularly when discrepancies in scientific nomenclature or taxonomic errors are present. We developed an R package, NTCmatch, designed to facilitate the identification of plant synonyms within the NTC list, enhancing the accuracy and efficiency of the ABS mechanism. The package was created using devtools and roxygen2, with readxl for data management. Results demonstrate the utility of NTCmatch in synonym matching, although the manual verification of plant parts and sources remains crucial. The package aims to streamline workflows for NBA, SBBs, and UTBCs, ensuring compliance with the Biological Diversity Act and facilitating the equitable sharing of benefits.

## INTRODUCTION

Equitable sharing of benefits arising from the commercial use of biological resources is one of the three foundational pillars of the Convention on Biological Diversity (CBD). This principle is crucial because it has the potential to drive the other two objectives of the CBD: the conservation of biological diversity and its sustainable use. As a signatory to the CBD, India has enacted the Biological Diversity Act in 2002 (amended in 2023) and the Biological Diversity Rules in 2004 to implement the provisions of the CBD within the country. The Act is implemented through a three-tier system, comprising the National Biodiversity Authority (NBA) at the central level, State Biodiversity Boards (SBBs)/Union Territory Biodiversity Councils (UTBCs) at the state/union territory level, and Biodiversity Management Committees at the local body level. The NBA, operating under the Ministry of Environment, Forests, and Climate Change, Government of India, is empowered to periodically issue guidelines related to the implementation of the Biological Diversity Act, 2002.

One of the critical regulatory frameworks under the Biological Diversity Act is the access to biological resources for commercial utilization, guided by the Nagoya Protocol of 2010 and subsequent guidelines issued by the NBA in 2014. These guidelines play a pivotal role in operationalizing the Access and Benefit Sharing (ABS) mechanism at both the central and state/UTBC levels. A key highlight of the guidelines is the establishment of operational slabs for calculating benefits based on the purchase price of biological resources or the gross ex-factory sale of the products derived from these resources.

Under the Biological Diversity Act, bioresources are defined as plants, animals, microorganisms, or parts thereof, including their genetic material and by-products (excluding value-added products) that have actual or potential use or value. However, this definition explicitly excludes human genetic material. Importantly, not all bioresources are subject to the ABS provisions of the Biological Diversity Act. Section 40 of the Act states: “Notwithstanding anything contained in this Act, the Central Government may, in consultation with the National Biodiversity Authority, by notification in the Official Gazette, declare that the provisions of this Act shall not apply to any items, including biological resources normally traded as commodities.”

The NBA and SBB/UTBC serve as the nodal organizations where applications for access to biological resources for commercial utilization can be submitted. This process requires the submission of Form I, as provided in the Biological Diversity Act, which serves as the primary document, supplemented by Form A, as notified in the ABS guidelines of 2014. Additionally, the submission must include the prescribed application fee and any other requisite documents. One of the crucial aspect of processing these forms at the NBA and SBB/UTBC level is the careful matching of the applied-for bioresources with the notified list of Normally Traded as Commodities (NTC). However, challenges arise when the scientific names in the application do not directly match those in the NTC list due to issues of synonymy. For instance, if an applicant refers to *Emblica officinalis*, they might not find this name on the NTC list because the list references *Phyllanthus emblica* instead. Similarly, the NTC list includes *Justicia adhatoda* but not its synonym, *Adhatoda zeylanica*. This discrepancy could lead to errors in processing if synonymy is not adequately considered. Further, there are typing errors in the notification viz. *Ailanthus excelsus* has been mentioned as *Ailanthus excelsa*, similarly *Anacardium occidentale* has been mentioned as *Anacardium occidendale* in the notification (NBA 2017). In addition, the compilation of the bio resource list for notification is an extensive task, leaving room for potential taxonomic errors.

To address these challenges, this work aims to develop an R package designed to search for synonyms of plants listed under the NTC list. This package will serve as a ready reference for the NBA, SBBs, and UTBCs working on Access and Benefit Sharing, thereby streamlining the workflow associated with processing Form I and Form A. The tool will enhance accuracy in the identification of bioresources, ensuring that applications are processed efficiently and in accordance with the correct taxonomic nomenclature.

## MATERIALS AND METHODS

### Data backbone

The notification of plants listed as normally traded as commodities was downloaded from the National Biodiversity Authority website (NBA 2016, 2017). The document in pdf format, was converted into MS Excel format using PDFgear application, allowing the extraction of the table listing the NTC names. A static copy of the taxonomic backbone data (version v.2023.12) was downloaded from worldflora website (WFO 2023). For the purpose of utilizing this data, WorldFlora R package (Kindt 2020) was installed. The WorldFlora package provides a straightforward pipeline for semi□automatic plant name checking, with success rates for credible name matches reported to range between 94.7% to 99.9% (Kindt 2020).

The scientific names listed under NTC were verified against the WorldFlora database and corrected where necessary. The comparison, was conducted using both worldflora website and WorldFlora R package. Any names that posed issues were addressed during this process.

Subsequently, a subset of names was stored separately from the main database. This subset included (i) fungi such as *Agaricus bisporus, Auricularia auricula-judae, Auricularia polytricha, Tremella fuciformis, Tremella mesenterica*, and *Pleurotus ostreatus*; and (ii) names with issues, such as those not listed in WorldFlora, including *Beta vulgaris* var. *cicla, Brassica campestris* var. *Brown sarson, Brassica campestris* var. *Yellow sarson, Dimorphotheca orientale, Dioscorea cayenensis* subsp. *rotundata, Eruca sativa* subsp. *sativa, Lilium asiaticum*, and *Pisum sativum* var. *arvense*, along with *Beta vulgaris* var. *bengalensis* to which WorldFlora package continuously applied fuzzy logic identification.

After the names were corrected, the list in the main database was searched for synonyms using the WorldFlora package. A final comprehensive list was compiled, incorporating the main database entries, their synonyms, and the previously separated fungal names and plant names not listed in WorldFlora.

### R package development

The R package development process utilized the *devtools* (Wickham et al. 2022) and *roxygen2* (Wickham et al. 2021) packages, with *readxl* (Wickham and Bryan 2023) as a dependency. The readxl package enables the reading of Excel files into a data frame in R, while the merge function is derived from base R. Since the readxl package may not be preinstalled on most machines, the following code is provided to facilitate its installation and loading into the R environment:

if(!require(readxl)){

install.packages(“readxl”)

library(readxl)}

### Package hosting and installation

NTCmatch package is hosted on github under the PradhanP-R repository (https://github.com/PradhanP-R/NTCmatch). It can be installed in R using the following commands:

install.packages(“remotes”)

remotes::install_github(“PradhanP-R/NTCmatch”)

## RESULTS AND DISCUSSION

The primary function of the NTCmatch package, match_NTC_names, is designed for synonym searching of plant names in the NTC list. This function requires the loading of user data, which is facilitated through the readxl package by reading data from an MS Excel spreadsheet into R. The user list should be formatted with an Excel sheet containing a column labeled ‘Name’ in cell A1, followed by the scientific names of plants without the author name, as depicted in Figure 1(A). The user list can be uploaded using the following command, which opens a graphical user interface for selecting the MS Excel file:

**Figure 1.**
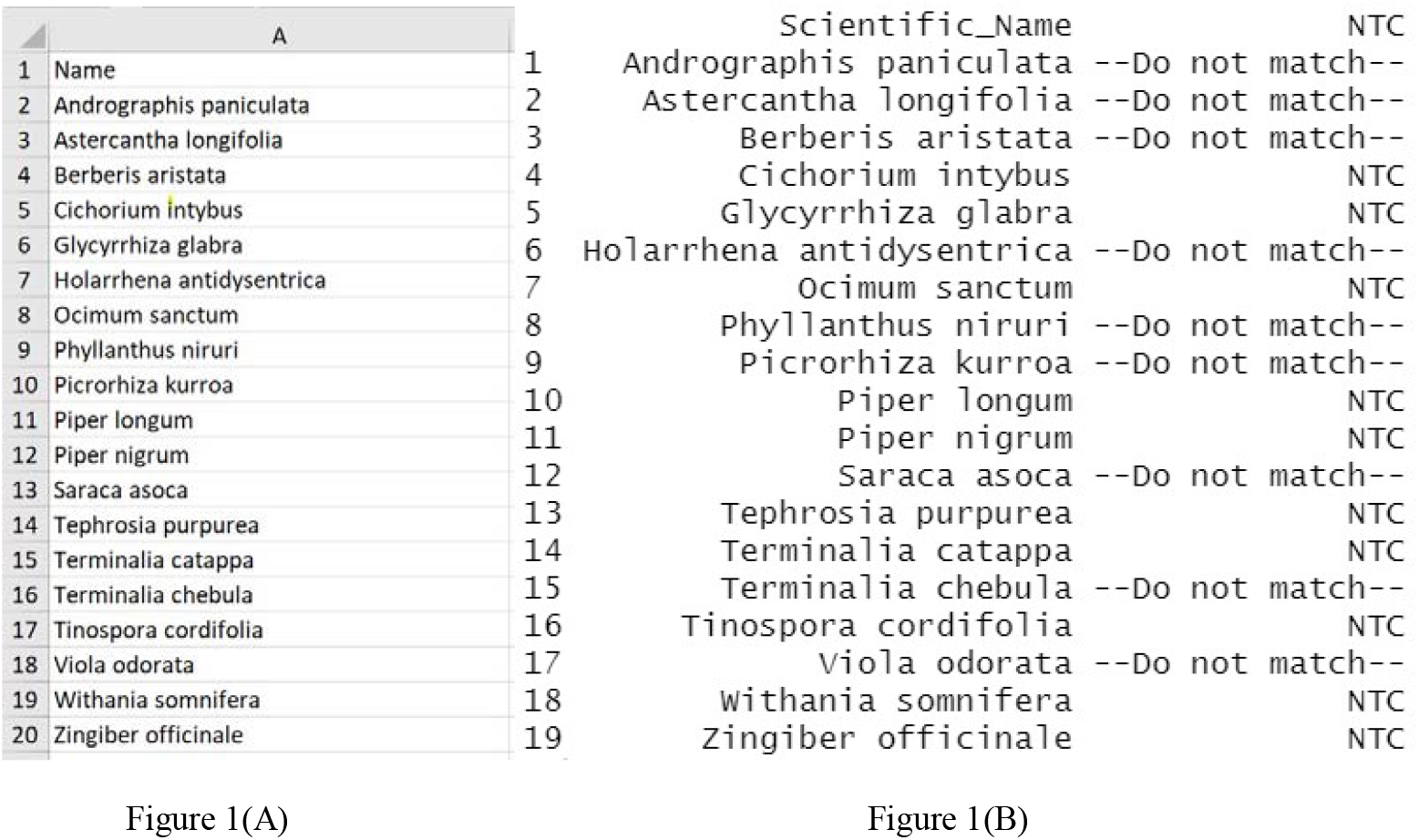
**A:** Demo user data with column with ‘Name’ head followed by scientific names, **B:** Output of function match_NTC_names() with the loaded user data

Userdata <- readxl::read_excel(file.choose())

Once the user data is loaded, the match results can be derived using the match_NTC_names function:

match_result <- match_NTC_names(Userdata)

The object names Userdata (containing the user list) and match_result (containing the result output) can be changed according to the user’s preference. However, the name of the object passed to the match_NTC_names() function must match the name of the object containing the user list.

The confirmation of the Normally Traded as Commodities (NTC) status of a biological resource involves a three-stage process. Firstly, the species name must be accurately matched. Secondly, the specific plant part applied for by the applicant must be identified and matched. Lastly, it is crucial to verify the source of the bioresource—whether it is derived from a cultivated source or a mixed source, including wild sources. While the R package being developed can significantly aid in identifying plant names and matching them with the NTC list, it is essential to note that the official processing Form I would still need to meticulously verify the plant part and the source to reach a definitive conclusion.

Incorporating the matching of plant parts directly into the NTCmatch package is indeed feasible and would streamline the process. However, this approach comes with a challenge. The NTC guidelines often list plant parts in singular or plural forms, while Form I applicants might describe plant parts in various ways. For instance, the NTC guidelines may list the plant part for *Cajanus cajan* as ‘Pod’. If the package includes both the singular ‘Pod’ and its plural ‘Pods,’ the search function would successfully match the NTC list if an applicant specifies ‘Pods’. However, if the applicant uses a different term, such as ‘fruit’ or ‘fruits,’ *Cajanus cajan* might not appear in the NTC list search results. Similarly, *Brassica oleracea* var. *gongylodes* is listed in the NTC guidelines with the plant part ‘Tuberous stem’. If the package accommodates variations like ‘Tuberous stem,’ ‘Tuberousstem,’ ‘Tuber,’ ‘Tubers,’ ‘Stem,’ and ‘Stems,’ these would likely match the NTC list. However, if an applicant describes the plant part using a term outside of these anticipated variations, the package would fail to recognize it as an NTC-listed resource.

Therefore, while the NTCmatch package can be enhanced to include plant part variations, there is an inherent limitation. The official processing the Form I must remain vigilant in manually cross-referencing the plant parts and sources mentioned by the applicant against the NTC list, ensuring no discrepancies occur due to variations in terminology. This careful attention to detail is critical to accurately determining the NTC status and ensuring compliance with the Access and Benefit Sharing (ABS) guidelines.

## CONCLUSION

The NTCmatch R package offers a significant tool for addressing the challenges in synonym matching of plant names listed under the Normally Traded as Commodities (NTC) within India’s Biological Diversity Act implementation framework. While the package streamlines the identification process and enhances the accuracy of applications under the Access and Benefit Sharing (ABS) mechanism, it has its limitations. The complexity of plant part nomenclature and the variability in applicant descriptions necessitate continued manual verification by officials. Incorporating plant part variations into the package is feasible but cannot eliminate the need for meticulous cross-referencing, especially regarding names of plant parts and source of bio resource. The development of NTCmatch represents a critical step toward improving the efficiency and effectiveness of ABS application processing, ensuring that the principles of equitable benefit sharing are upheld in line with the objectives of the Convention on Biological Diversity. Future improvements could focus on expanding the package’s functionality to better accommodate the nuances of plant part identification, further reducing the potential for errors in processing Form I applications.

## Abbreviations (if any)

ABS: Access and Benefit Sharing
CBD: Convention of Biological Diversity
NBA: National Biodiversity Authority
NTC: Normally Traded as Commodities
SBBs: State Biodiversity Boards
UTBCs: Union Territory Biodiversity Councils

